# Neural correlates of masked and unmasked tones: psychoacoustics and late auditory evoked potentials (LAEPs)

**DOI:** 10.1101/2021.11.06.467541

**Authors:** Hyojin Kim, Bastian Epp

## Abstract

Hearing thresholds are commonly used to quantify a listener’s ability to detect sound. In the presence of masking sounds, hearing thresholds can vary depending on the signal properties of the target and the masker, commonly referred to as auditory cues. Target detection can be facilitated with comodulated masking noise and interaural phase disparity (IPD). This can be quantified with a decrease in detection thresholds or masking release: comodulation masking release (CMR, for comodulation) and binaural masking level difference (BMLD, for IPD). As these measures only reflect the low limit of levels for target detection, the relevance of masking release at supra-threshold levels is still unclear. Here, we used psychoacoustic and electrophysiological measures to investigate the effect of masking release for a masked tone at supra-threshold levels. Behaviorally, we investigated how the amount of masking release affects the salience at supra-threshold levels. We used intensity just-noticeable difference (JND) to quantify level-dependent changes in the salience of the tonal signal. As a physiological correlate, we investigated late auditory evoked potentials (LAEPs) with electroencephalography (EEG). The results showed that the intensity JNDs were equal at the same physical target tone level, regardless of the presence or absence of masking release. Estimated salience was correlated with the amount of masking release. However, salience measures across conditions converged with the target tone level above 70 dB SPL. For the LAEPs, the P2 amplitudes were more closely linked to behavioral measures than the N1 amplitudes. Both behavioral and electrophysiological measures suggest that the salience of a masked tone at supra-threshold levels is correlated with the amount of masking release.

## 1. Introduction

Acoustic scenes in everyday life consist of a complex mixture of sounds. Our auditory system can segregate this mixture into a target sound and a noise background, enabling communication in acoustically complex environments. It is assumed that our auditory system binds various acoustic features arising from the same source into a sound object or an acoustic stream. As an example of such features, speech shows coherent amplitude modulation patterns across a wide frequency range (Raphael et al., 2007). Previous studies have shown that coherent modulation, or comodulation, is beneficial for the detection of a tone in noise (Hall et al., 1984; Nelken et al., 1999). This suggests that comodulation can be used to group spectral components across a wide range of frequency bands as comodulation indicates that these components likely stem from the same source. Such grouping can facilitate the segregation of the target signal from the noise and result in an enhanced target detection. Similarly, spatial information can also facilitate sound detection. When an acoustic source is lateralized relative to the listeners’ head direction, interaural disparities between the ears can be induced. For instance, an interaural phase difference (IPD) can facilitate the target identification by grouping acoustic features from the same source. For instance, when the target tone in the noise is presented with an IPD between left and right ears, detection thresholds of the tone are lower compared to the case with no IPD (van de Par and Kohlrausch, 1999).

In psychoacoustics, such enhancement in detection performance is considered as “release from masking,” and referred to as masking release. Masking release can be quantified as the amount of decrease in the detection threshold in the presence of beneficial cues compared to the detection threshold obtained in the absence of these cues. A decrease in detection threshold by comodulation is termed as comodulation masking release (CMR), and the decrease related to a binaural cue is referred to as binaural masking level difference (BMLD). In simple cases where the target tone is presented with both comodulation and IPD cue, the amount of masking release was found to be a superposition of CMR and BMLD (e.g., Epp and Verhey, 2009). In their study, they interpreted the psychoacoustical measures of CMR and BMLD as the result of enhanced neural representations by bottom-up, serial neural processing. A reduced CMR in the presence of BMLD was found by Hall III et al. (2011), indicating a small interaction between the processes underlying CMR and BMLD. For CMR, the earliest physiological neural correlate of CMR was found at the CN level (Pressnitzer et al., 2001; Neuert et al., 2004). The neuronal representation of the comodulated signal gets sharper at the inferior colliculus (IC) level (Nelken et al., 1999). For BMLD, neural correlates of IPD were found at the IC level (Shackleton et al., 2003, 2005; Zohar et al., 2011). Based on these findings, Epp and Verhey (2009) suggested that the superposition of CMR and BMLD is the combination of the enhanced internal signal-to-noise ratio of the neural representations.

When a comodulated masker is preceded and followed by another masker (temporal fringe), CMR can be reduced or increased depending on the preceding and following maskers (Grose et al., 2009; Dau et al., 2009, 2005). It is not yet clear whether this superposition of CMR and BMLD also applies in cases where the amount of CMR is affected by the temporal context of the spectral masker components. When the temporal fringe is comodulated, CMR can be enhanced compared to the absence of the fringe (Grose et al., 2009). One interpretation of this result is that the fringe facilitates the grouping of the masker, thereby enhancing the separation of the target sound from noise. On the other hand, when the temporal fringe is uncorrelated, CMR can be reduced compared to a condition without temporal fringe (Grose et al., 2009). In this case, the fringe has a detrimental effect on the grouping of the masker, resulting in reduced CMR. Hence, these results can be linked to stream formation by frequency grouping in time, suggesting the influence of the high-level auditory processing on CMR (Grose et al., 2009; Dau et al., 2005, 2009). Neural correlates of the effect of preceding maskers on target detection were found at the cortical level (A1) (Sollini and Chadderton, 2016). In their study, neural responses to the stimuli were enhanced by a preceding comodulated masker compared to the ones preceded by an uncorrelated masker. Nevertheless, it remains unclear whether the improved neural response to the target tone at the A1 level is the result of relayed encoding from the CN to A1, or an additional encoding at the A1 level, or a cortical feedback from A1 to CN (Sollini and Chadderton, 2016). Furthermore, little is known about the effect of stream formation on the masking release induced by interaural disparities like BMLD.

CMR and BMLD are well characterized at low intensities (near thresholds). However, one might argue that target detection in communication occurs at levels well above threshold, i.e., supra-threshold levels. This leads to a question of the relevance of CMR and BMLD to communication in complex acoustic environments. Several studies have investigated the perception at supra-threshold levels in masking release conditions. A common goal was to map physical properties (e.g., the increment in the intensity of a sound) to psychophysical variables (e.g., the increment in loudness or salience). Related studies used categorical loudness scaling (Verhey and Heeren, 2015) and continuous scaling of the perceived salience (Egger et al., 2019). In categorical loudness scaling, Verhey and Heeren (2015) used a matching method where listeners matched the loudness between the target tone in modulated noise and in unmodulated noise. The level of unmodulated noise was reduced by the amount of threshold difference between the two noise types. Their results showed that the supra-threshold perception of the target tone in the modulated noise was similar to that in the unmodulated noise at a reduced level, suggesting that the masking release results from reduced internal masker level. With continuous scaling tested on both CMR and BMLD, the data from Egger et al. (2019) showed individual variability in the ratings, presumably because some listeners confused the loudness of the overall sound with the salience of the target tone in noise. The limitation of those methods is that those measures strongly depend on listeners’ subjective criteria for decision-making. As an alternative to asking listeners to quantify the salience, we used a just-noticeable difference (JND). With this method, the intensity of a target signal is decreased step-wise until the listener cannot detect the change in intensity relative to the reference signal with a fixed level (intensity JND). This approach might potentially reduce the impact of subjective criteria for judging the salience of the target tone. A previous study showed that the intensity JND follows the power law (Ozimek and Zwislocki, 1996). However, whether this relationship holds for the salience in masking release conditions is unclear.

As a neural correlate of an enhanced internal representation of the target tone in noise, Epp et al. (2013) evaluated auditory evoked potentials (AEPs). They hypothesized that that the neural representation of the target tone in masker at the cortical level is correlated with the level above masked threshold. They assumed that the physical signal-to-noise ratio of the masked tone is enhanced by comodulation and IPD cues along the auditory pathway. The resulting enhanced neural representation of masked tone would be reflected in the peaks of the AEPs. They measured AEPs at various intensities of the tonal component with a fixed masker level. They found that the amplitude of the P2 component of the late auditory evoked potentials (LAEPs) was proportional to the amount of masking release, CMR, and BMLD. The growth function of the P2 amplitude was similar across conditions. Based on this finding, the follow-up study by Egger et al. (2019) hypothesized that LAEPs measured at the same level above masked threshold (e.g., threshold + 5 dB, + 10 dB, etc.) will evoke the same amplitude of the P2 components regardless of masking release conditions. They measured LAEPs at the six different supra-threshold levels, together with the salience of the tonal component in maskers with the scaling method. The results of LAEPs showed that the tone at the same supra-threshold levels evoked similar P2 amplitudes. However, ratings of the salience were not correlated with P2.

To shed light on the mechanism and neural representation of CMR and BMLD, we investigated: i) the effect of stream formation induced by preceding maskers on CMR and BMLD. We hypothesized that if stream formation results from high-level auditory processing (e.g., integration of temporal context across frequencies), both CMR and BMLD will be affected by stream formation. Understanding the interaction of bottom-up processing and the stream formation will help to reveal the neural encoding strategies underlying sound source separation. ii) the intensity JND as a measure of the “internal representation” of the tonal component in masking release conditions at supra-threshold levels. As an extension of Ozimek and Zwislocki (1996) and Egger et al. (2019), we hypothesized that the intensity JND in masking release would follow a power law as a function of supra-threshold levels, regardless of masking release conditions. If the internal neural representation is enhanced proportional to the amount of masking release, the condition with a higher amount of CMR and BMLD will show lower intensity JNDs at the same physical target tone level. iii) the correlation between the intensity JND and LAEP measures. In the present study, we estimated the slope of changes in P2 amplitudes with increased levels. We hypothesized that the increment in P2 with increasing tone level would be inversely proportional to the intensity JND. In addition, based on the intensity JND measures, we estimated the salience and investigated whether P2 can reflect the estimated salience behaviorally.

## 2. Materials and methods

### 2.1. Stimuli

Our study consisted of three experiments: i) psychoacoustical threshold measurements to quantify CMR and BMLD; ii) intensity JND measurements; iii) EEG experiments for measuring LAEPs. For all three experiments, we used the same eight masking release conditions (Fig 1). The stimulus consisted of five noise bands as a masker and a pure tone as a target signal: one noise band was centered at the frequency of the target tone (center band, CB). The other bands were equally spaced with a distance of 120 Hz above and below the CB (flanking bands, FBs). Each masker band had a bandwidth of 20 Hz and a level of 60 dB SPL. The target tone was centered at 700 Hz. We chose this frequency setting to maximize the effect of the stream formation on CMR based on the previous work by Grose et al. (2009). Each interval consisted of a preceding masker with a duration of 500ms (“preceding masker”) and a masked target tone with a duration of 200ms (“masked tone interval”). We used four masker conditions. In the reference condition, the maskers had random intensity fluctuations across frequency for both the “preceding masker” and the “masked tone interval” (RR). In the three other conditions, the target tone was embedded in comodulated noise and preceded by one of the following masker types: a masker with random intensity fluctuations across frequency (RC), a comodulated masker (CC), and a masker where only the FBs were comodulated (FC). All maskers were presented diotically and had 20 ms raised-cosine on- and offset ramps. For RC and FC conditions, the same on- and offset ramps were added in the transition with a 50% overlap. The noise bands were generated in the frequency domain and transformed into the time domain. The noise bands were assigned numbers from a uniformly distributed random process to the real and imaginary parts of the respective frequency components. For the R masker, different numbers were assigned for each noise band. For the C masker, the same numbers were used for all five noise bands. The stimuli were generated with newly drawn numbers for each interval and each trial. To induce BMLD, the target tone was presented with an IPD of 0 or *π* in combination with the same four masker types, leading to a total of eight stimulus conditions.

**Fig 1:**
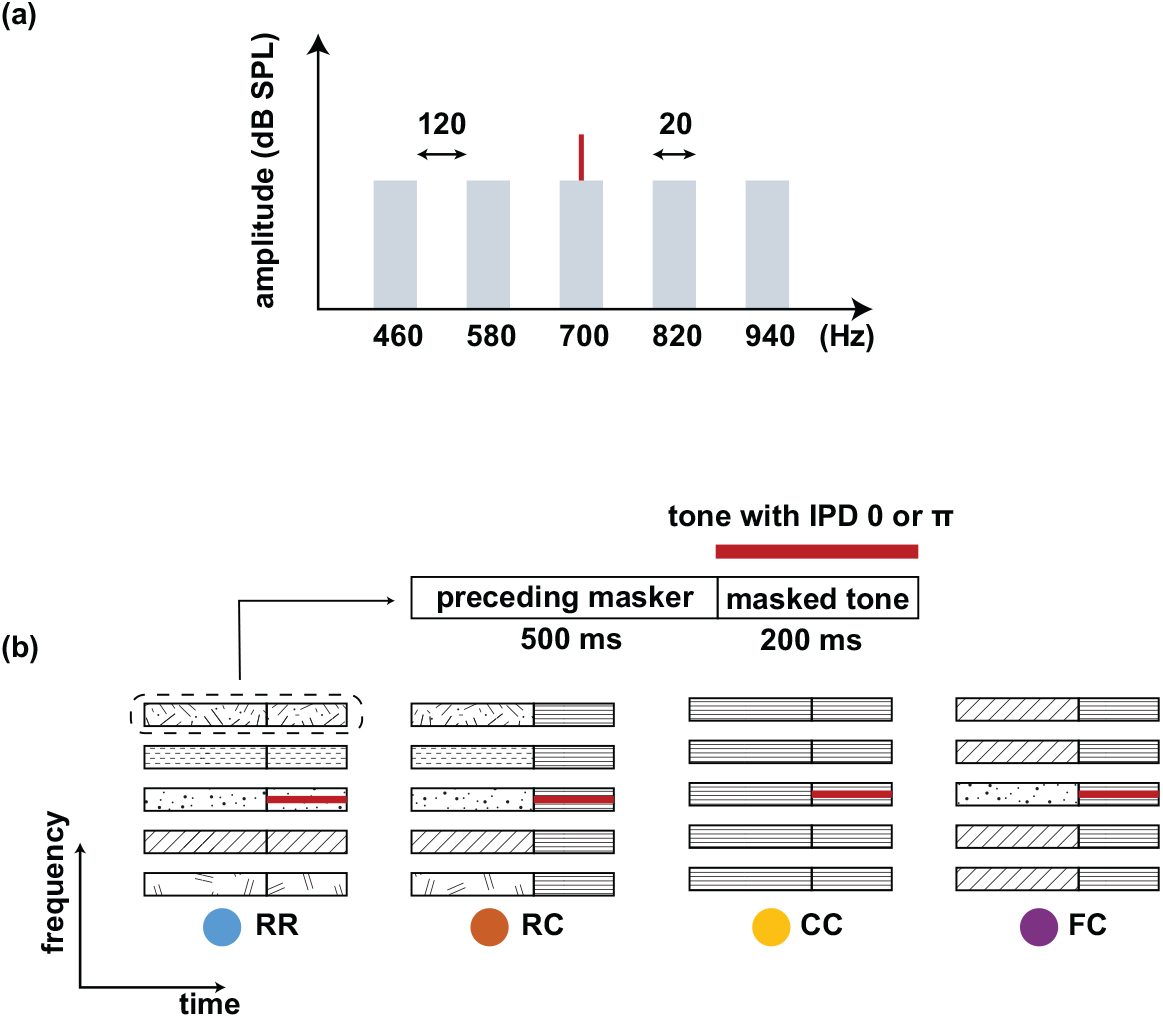
**(a) Spectra of the stimulus. A target tone (700 Hz) was presented with a masking noise consisting of five narrow-band maskers: One band centered at the target tone frequency (center band, CB) and four flanking bands (FBs). The bandwidth of each masker band was 20 Hz, and the frequency spacing between FBs was 120 Hz. The overall level of the noise was set to 60 dB SPL. (b) Schematic spectrograms of the stimuli. Each stimulus consists of a preceding masker (500 ms) and masked tone (200 ms). Four types of maskers were used: RR, RC, CC, and FC. The RR was used as the reference condition with uncorrelated masker bands. In the other three conditions, the maskers consisted of a comodulated masker preceded by three different maskers: uncorrelated masker (RC), comodulated masker (CC), and the masker with comodulated flanking-bands (FC). The thick red line represents a tone that was presented with an IPD of 0 or** *π*.

### 2.2. Apparatus

During all three experiments, the listeners were seated in a double-walled, soundproof booth. All stimuli were generated in MATLAB 2018b (TheMathworks, Natick, MA) with a sampling rate of 44100 Hz and a 16-bit resolution, converted from digital to analog (RME Frieface UCX), amplified (Phonitor mini, SPL electronics), and played back through headphones (ER-2, Etymotic Research). The headphones were calibrated at the signal frequency of the tone. For the recording of AEPs, we used a g.Tec HIamp system with a sampling rate of 1024 Hz. The 64 channels of active electrodes were set up with highly conductive electrode gel to reduce the impedance between the scalp and electrodes. The reference electrodes were placed close to the mastoid of both ears and the other electrodes were placed based on g.GAMMAcap 64 channel setup from g.Tec.

### 2.3. Listeners

We recruited fifteen normal-hearing listeners. None of them reported any history of hearing impairment. All but one listener had pure-tone hearing thresholds within 15 dB HL for the standard audiometric frequencies from 125 to 4000 Hz. One listener was tested with 20 dB at 125 Hz. All participants provided informed consent, and all experiments were approved by the Science-Ethics Committee for the Capital Region of Denmark (reference H-16036391). All of them participated in the first experiment, eleven of them participated in the second experiment, and ten of them participated in the third experiment.

### 2.4. Procedure

In the first experiment, we measured masked thresholds individually for the eight stimulus conditions presented in random order. We used an adaptive, three-interval, three-alternative forced-choice procedure (3-AFC) with a one-up, two-down rule to estimate the 70.7% of the psychometric function (Ewert, 2013; Levitt, 1971). Two intervals contained the masking noise only. The remaining interval contained the target tone in addition to the masker. The three intervals were presented with a temporal gap of 500 ms in between. The listeners’ task was to select the interval with the target tone by pressing the corresponding number key (1, 2, 3) on the keyboard. Visual feedback was provided, indicating whether the answer was “WRONG” or “CORRECT”. The initial level of the target tone was set to 75 dB SPL and was adjusted with an initial step size of 8 dB. The step size was halved after each lower reversal until it reached the minimum step size of 1 dB. The signal level at a minimum step size of 1 dB was measured six times, and the mean of the last six reversals was used as the estimated threshold. Each listener performed three threshold measurements for all conditions. The average of three measurements was used as individual masked thresholds for the next two experiments. Additional measurements were performed if the thresholds from the last three measurements had a standard deviation larger than 3 dB.

In the second experiment, we measured intensity JNDs individually at six supra-threshold levels for all conditions. The intensity of the tone was individually adjusted for each listener to match levels of +0 dB (threshold), +5 dB, +10 dB, +15 dB, +20 dB, and +25 dB relative to the threshold. The individual mean of three threshold measurements from the first experiment was used to set the reference of +0 dB. We used the same setup and 3-AFC method as for the first experiment. Two intervals contained the masked target tone with a fixed level at one of the supra-threshold levels (“reference interval”), and the remaining interval contained the masked target tone with a higher level than the others (“target interval”). The intervals were presented with a temporal gap of 500 ms in between. Listeners were asked to select the interval with the tone of highest intensity by pressing the corresponding number key (1, 2, 3) on the keyboard. Visual feedback was provided, indicating whether the answer was “WRONG” or “CORRECT.” The order of conditions and supra-threshold levels were randomized. The initial level of the tone in the target interval was set to 75 dB SPL. The level of the target tone was adjusted with the initial step size of 8 dB. The step size was halved after each lower reversal until it reached the minimum step size of 1 dB. The signal level at a minimum step size of 1 dB was measured six times, and the mean of the last six reversals was used as the JND. Listeners were familiarized with the task by a test run. Each listener performed three trials for all conditions. If the supra-threshold level exceeded 80 dB, the intensity JND measure was skipped. We calculated the intensity JND by subtracting the level of “reference intervals” from the minimum level of discriminable tone in “the target interval”.

In the third experiment, we measured late auditory evoked potentials (LAEPs) at three supra-threshold levels for all conditions. The intensity of the tone was individually adjusted for each listener to match levels of +15 dB, +20 dB, and +25 dB above the threshold. The individual mean of three threshold measurements from the first experiment was used to set supra-threshold levels. The stimuli for each condition and supra-threshold level were presented 400 times in random order. In addition, noise-only stimuli were presented 40 times for each condition. The presentations were separated by a random inter-stimulus interval of 500ms with jitter. During the experiment, a silent movie with subtitles was presented on a low-radiation screen. The listeners were asked to sit comfortably and avoid movement as much as possible. The experiment was divided into six blocks of approximately 38 minutes each. These were divided into two sessions on different days.

## Data analysis

### The threshold measurements

We used CMR and BMLD to quantify the amount of masking release in eight conditions. We used several acronyms for masking release measures for each condition as follows. For comodulation masking release (CMR),

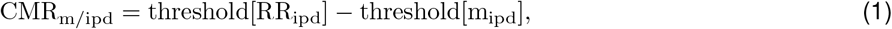

Here, *m* stands for one of three masker types (RC, CC, FC) and *ipd* stands for the IPD of the tone between two ears (0 or *π*). As an example, 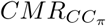 is the amount of a decrease in threshold in *CC*_*π*_ condition compared to *RR*_*π*_ condition. A positive value indicates a decreased detection threshold, and a negative value indicates a increased detection threshold. For binaural masking level difference (BMLD),

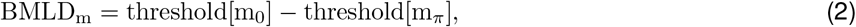

Here, *m* stands for one of three masker types (RC, CC, FC). As an example, *BMLD*_*CC*_ is the amount of a decrease in threshold in *CC* condition with IPD of *π* compared to *CC* condition without IPD. For statistical analysis, the Lilliefors test was used for a normality test. To compare CMR and BMLD across four masker types, one-way ANOVA followed by Tukey’s multiple comparison tests were used. In the case where the data did not follow a normal distribution, the Kruskal–Wallis test was used, followed by Dunn’s multiple comparison test. To compare CMR between two conditions with the same masker type but with different IPD, Wilcoxon signed-rank test was used.

### Intensity JNDs

We calculated the intensity JND by subtracting the intensity level of the reference intervals from the minimum intensity level of the discriminable tone in the target interval (Δ*L*). We fitted JND measures for each condition with a power law. In addition, we estimated the Weber fraction *k* by dividing the intensity JND (Δ*L*) with the intensity level of the target tone (*L*). We also calculated 10*log*(Δ*L/L*). From the fitted intensity JNDs, we estimated the salience from 1 to 10 (arbitrary scale). For each condition, we assumed that the salience was one at the corresponding masked thresholds. We increased the salience by one when the level was increased by the intensity JND at the current level. We repeated this estimation until the salience reached ten.

### Late auditory evoked potentials (LAEPs)

Collected data were analyzed using FieldTrip (Oostenveld et al., 2011). In short, the EEG data were partitioned into epochs from -300 to 850 ms relative to the onset of the preceding masker. The region of interest was the central position (Cz), and the reference signals were the average of two electrodes near the mastoids. Each epoch was low-pass (Butterworth IIR filter, 6th order, zero-phase) filtered with a cut-off frequency of 20 Hz. Detrending, baseline correction, and weighted averaging were applied to increase the signal-to-noise ratio (Riedel et al., 2001). Trials containing signals exceeding 100 *µ*V in any channel were rejected as artifacts. For auditory evoked potentials (AEPs), we extracted the signals from 100 ms before the onset of the target tone and 100 ms after the offset of the target tone from the averaged epochs. Baseline correction was applied considering a 100 ms pre-stimulus period. The grand mean of AEPs was computed with arithmetic mean over all individual AEPs. We selected the first negative component (N1) and the second positive component (P2) as a peak measure individually. We defined the peak of the first negative deflection in the time window between 100 ms and 200 ms (with respect to the target onset) as N1 and the peak of the second positive deflection in the time window between 200 ms and 300 ms as P2. This was estimated for each individual AEPs to eliminate individual differences in latency. Peak amplitudes were extracted by the MATLAB function *findpeaks* by locating minima and maxima within the time frame defined for N1 and P2, respectively. Extracted LAEPs were visually verified. In the case where multiple components were found, the one with the largest amplitude was selected. When there was no component found, this condition was excluded from the analysis.

## 3. Results

### 3.1. Experiment 1. Masked thresholds

Fig 2a shows the mean masked thresholds for eight stimulus conditions. For an IPD of 0, thresholds were highest for the FC condition and lowest for the CC condition. The observed mean threshold across all the participants for the *RR*_0_ condition was 55.4 dB. The *RC*_0_ condition had a mean threshold of 52.2 dB. In the *CC*_0_ condition, the threshold was 45.7 dB. In the *FC*_0_ condition, the mean threshold was found to be 58.7 dB. The same overall pattern of the thresholds was found for an IPD of *π* with the highest threshold for the FC condition and the lowest for the CC condition. The *RR*_*π*_ had a mean threshold of 39.5 dB, and the *RC*_*π*_ condition had a mean threshold of 38.3 dB. In the *CC*_*π*_, the mean threshold was 33.1 dB, and that of the *FC*_*π*_ condition was 45.7 dB.

**Fig 2:**
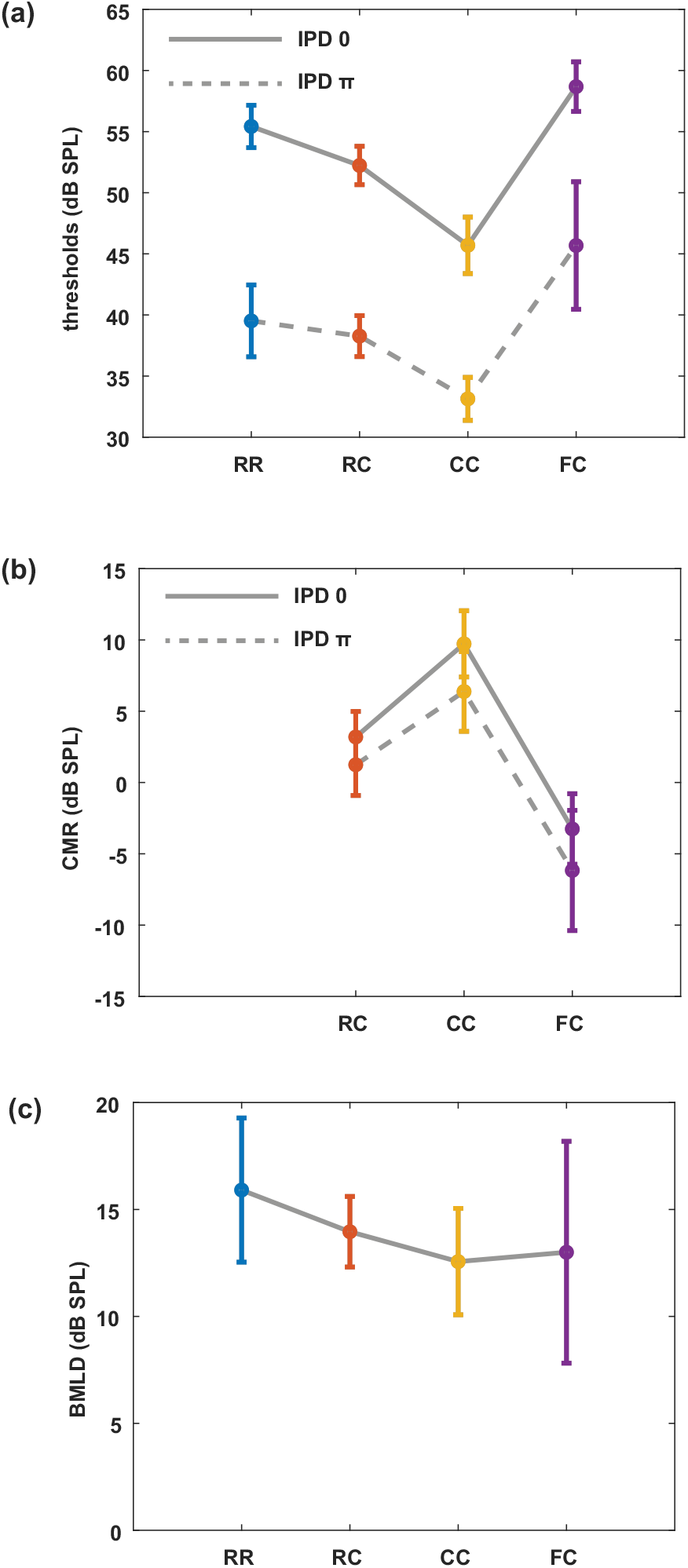
**Mean masked thresholds for all conditions and masking releases. (a) Masked thresholds from eight masking release conditions averaged over all listeners. (b) CMR with the RR masker as a reference. (c) BMLD for all masker types. Error bars indicate plus-minus one standard deviation**.

Fig 2b shows the CMR calculated for each condition by using the RR condition as reference (eq. (1)). The CMR was highest in CC conditions. While the CMR were positive for RC and CC conditions, FC conditions showed negative CMR. In the diotic conditions, *CMR*_*RC*0_ was 3.2 dB, *CMR*_*CC*0_ was 9.7 dB and *CMR*_*FC*0_ was -3.3 dB. In dichotic conditions, *CMR*_*RCπ*_ was 1.2 dB, *CMR*_*CCπ*_ was 6.4 dB and *CMR*_*FCπ*_ was -6.2 dB. Statistical analysis showed that CMR measures were different between different masker types. In diotic conditions, there was a significant difference in CMR between masker types (Kruskal-Wallis, p<0.05). Likewise, in dichotic conditions, CMR was significantly different between masker types (Kruskal-Wallis, p<0.05). Between diotic and dichotic conditions with the same masker type, all masker types showed a significant difference (Wilcoxon signed-rank test, p<0.05). Fig 2c shows the BMLD calculated for each condition by using the threshold in the corresponding diotic condition as reference (eq. (2)). *BMLD*_*RR*_ was 15.9 dB, *BMLD*_*RC*_ was 13.9 dB, *BMLD*_*CC*_ was 12.6 dB, and *BMLD*_*FC*_ was 13 dB. Multiple comparison tests showed that the *BMLD*_*RR*_ and *BMLD*_*CC*_ were significantly different (one-way ANOVA, p<0.05).

### 3.2. Experiment 2. Intensity JNDs

Fig 3 shows the individual intensity JND measures as the function of the physical target tone level in the reference signal. Each panel shows the intensity JND measures at supra-threshold levels ranging from threshold (+ 0 dB) to + 25 dB in four masker types with both IPD of 0 (solid line) and IPD of *π* (dashed line). For each masker type, the intensity JND measures were fitted with a power function. Additionally, the intensity JND measures (Δ*L*) were re-scaled as 10*log*(Δ*L/L*), and fitted with a power function (Fig 4). Overall, conditions with lower detection thresholds (e.g., *CC*_*π*_) showed higher JNDs compared to those with higher detection thresholds (e.g., *RR*_0_). This indicates that the degree of enhancement in the salience depends on the target tone level rather than the supra-threshold level. Fig 5 shows the averaged intensity JND (left) and re-scaled JND (right) as the function of the physical target tone level in the reference signal. The intensity JND measures of all conditions and listeners are shown with scatter plots and fitted with the power function. The intensity JNDs decreased with an increasing level of the target tone in all masking release conditions. Re-scaled JND measures showed better goodness of fit with the power function.

**Fig 3:**
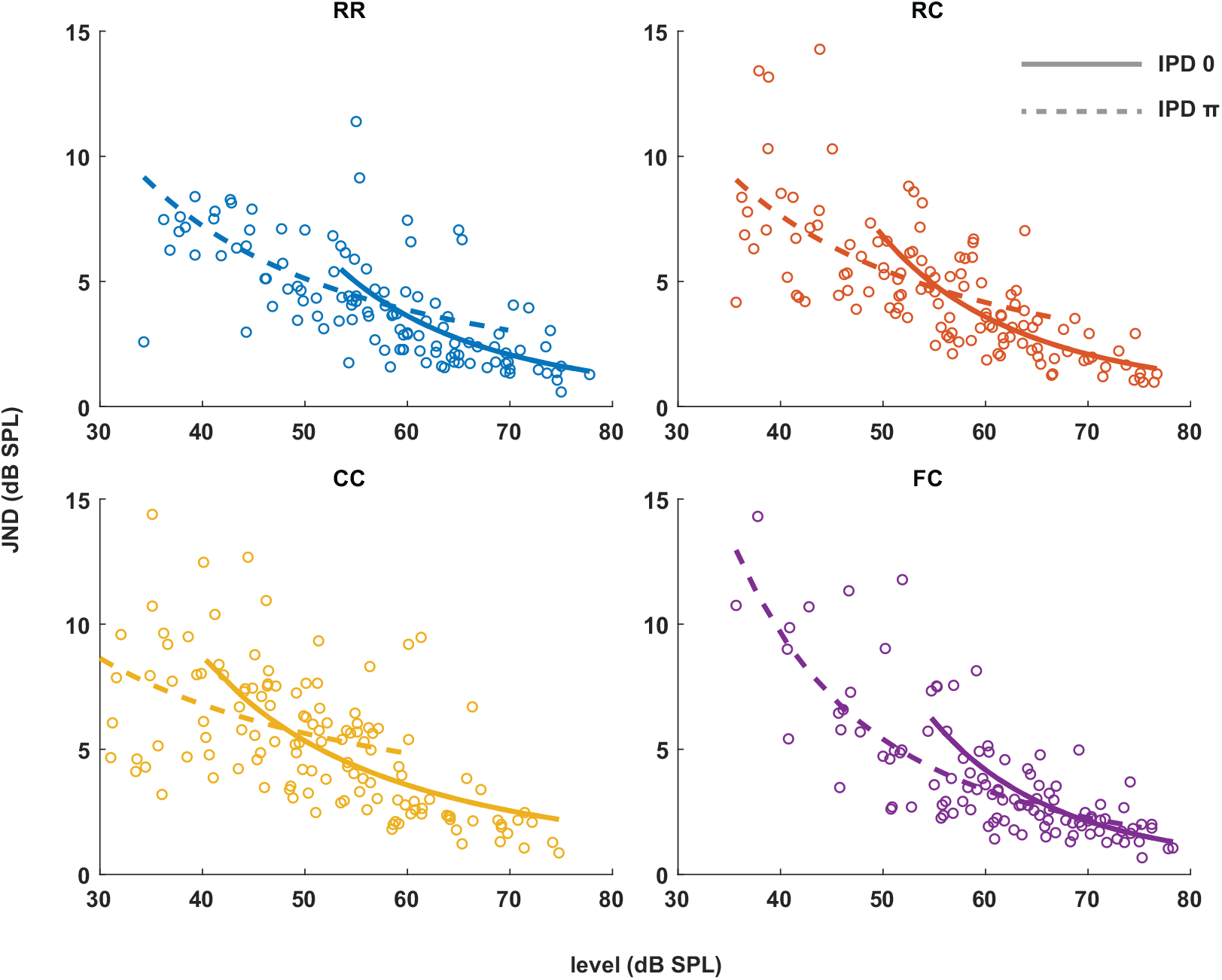
**Intensity JNDs for all stimulus conditions and power law fit for diotic (solid line) and dichotic target signal (dashed line). Individual data are plotted as single points. The data for each condition are fitted with a power function. Each color represents four masker types. Solid lines represent diotic conditions (IPD of 0) and dotted lines represent dichotic conditions (IPD of** *π***). The goodness of fit for each condition with IPD of 0 was: RR(***R*^2^**=0.4748), RC(***R*^2^**=0.6614), CC(***R*^2^**=0.4732), FC(***R*^2^**=0.5604). The goodness of fit for each condition with IPD of** *π* **was: RR(***R*^2^**=0.2743), RC(***R*^2^**=0.1613), CC(***R*^2^**=0.2946), FC(***R*^2^**=0.6170)**.

**Fig 4:**
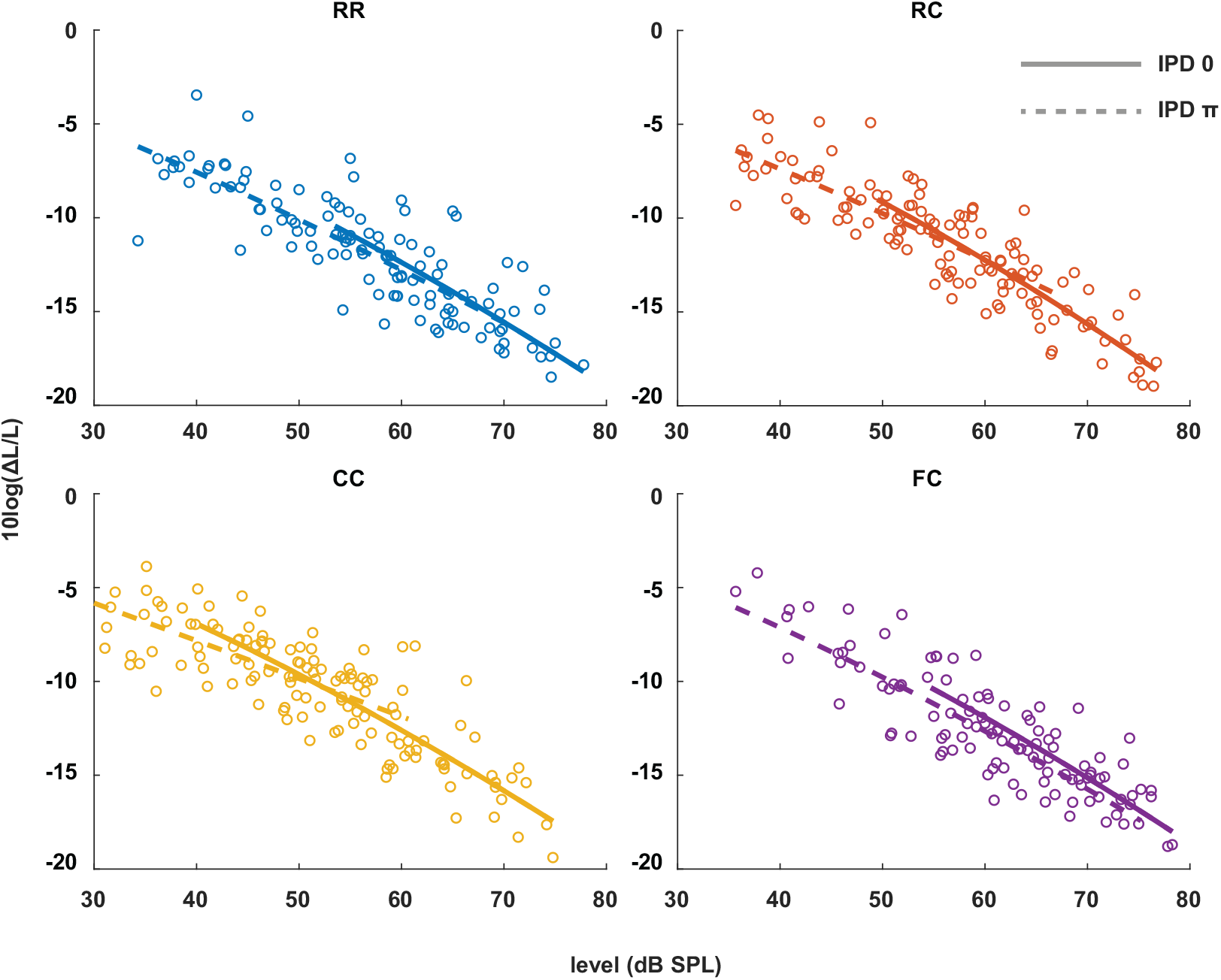
**Re-scaled intensity JND measures. Individual data are plotted as single points. The data for each condition are fitted with a power function in the same manner as the Fig 3. The goodness of fit for each condition with IPD of 0 was: RR(***R*^2^**=0.6386), RC(***R*^2^**=0.7914), CC(***R*^2^**=0.7472), FC(***R*^2^**=0.6491). The goodness of fit for each condition with IPD of** *π* **was: RR(***R*^2^**=0.6338), RC(***R*^2^**=0.6316), CC(***R*^2^**=0.4993), FC(***R*^2^**=0.7559)**.

**Fig 5:**
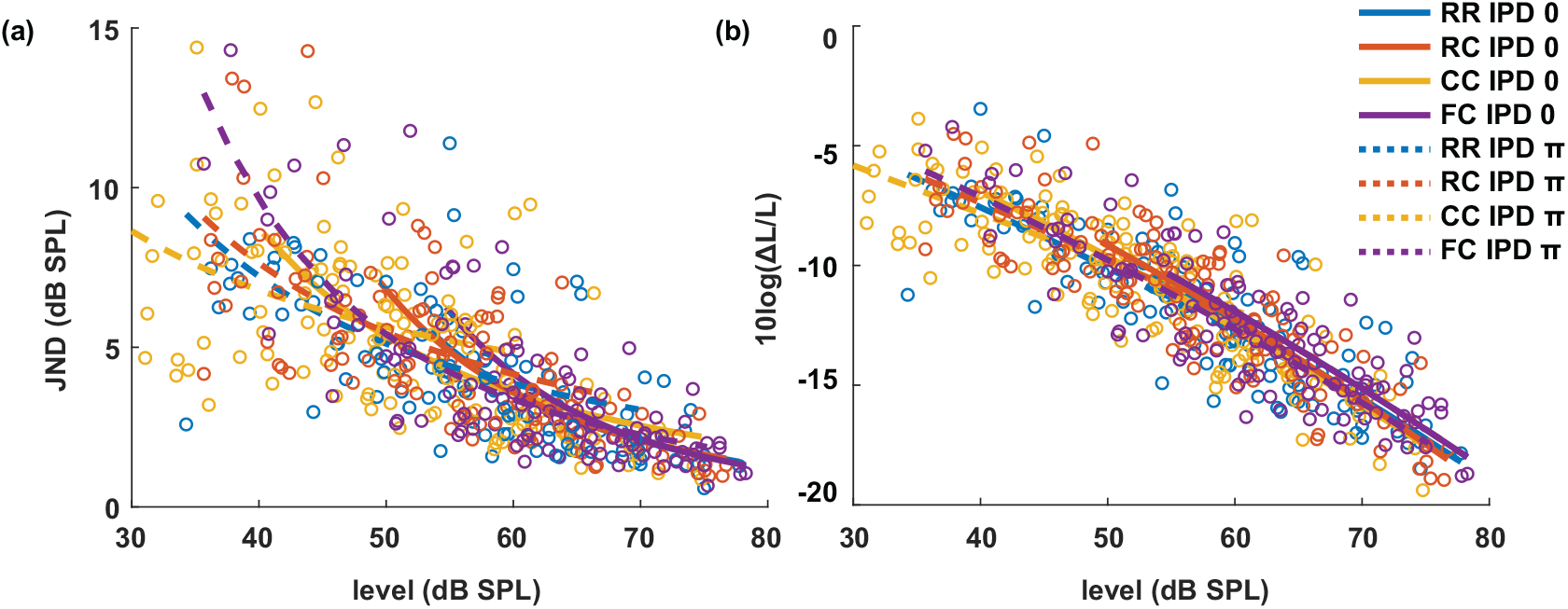
(a) The intensity JND measures. (b) Re-scaled intensity JND measures across all conditions.

#### Estimated salience

We define salience in the context of this study as the perceptual quantity that describes how clear the tone is perceived in noise. Assuming that the salience is the same as one at the threshold level, the salience would increase as the level of the target tone is increased. We hypothesized that the salience rating would increase by one the target tone level is increased by the intensity JND. The estimated salience is shown in Fig 6. At the same physical target tone level, estimated salience was higher for conditions with lower detection thresholds. For instance, the salience was higher in dichotic conditions compared to the diotic conditions with the same masker type. Estimated salience converged when the target tone level was above around 70 dB SPL.

**Fig 6:**
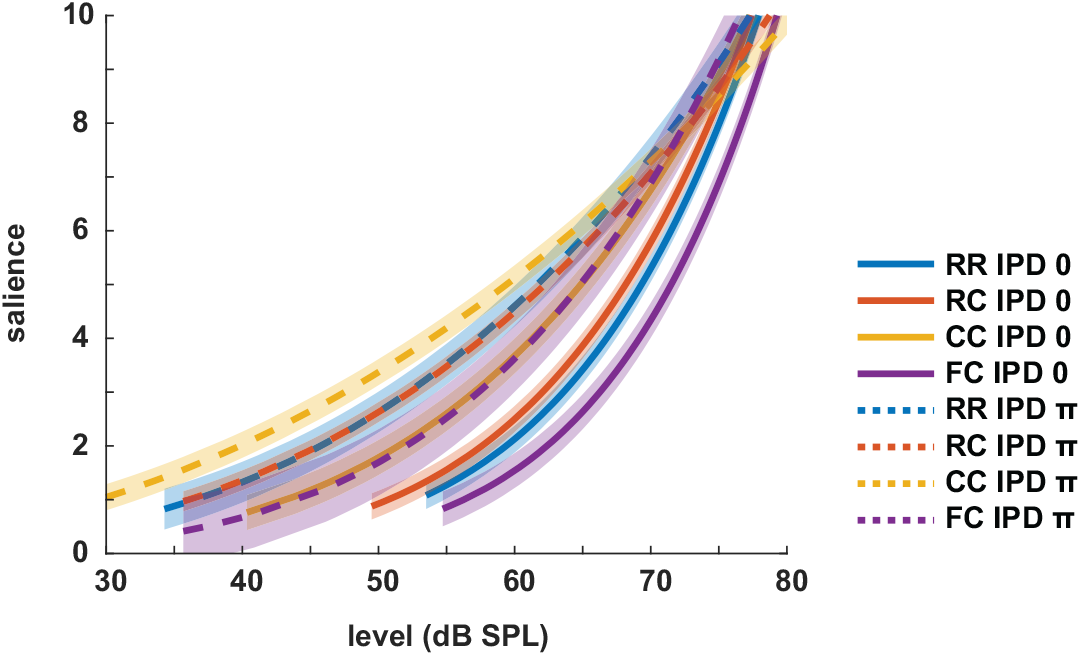
**Estimated salience ratings. Each color represents four masker types. Solid lines represent diotic conditions (IPD of 0) and dotted lines represent dichotic conditions (IPD of** *π***). The shaded areas indicate +/- one root mean square error**.

### 3.3. Experiment 3. Late auditory evoked potentials

Fig 7a shows the grand mean AEPs across all listeners for each condition. The plot shows the AEPs to diotic signals (solid lines) and dichotic signals (dashed lines) in the four maker types *RR, RC, CC*, and *FC*, respectively. Following the onset of the stimuli (Fig 7a, blue line), an onset response was elicited, which went back to a constant value after around 300 ms post-onset. The response to the target signal was found from around t=550 ms. Fig 7b shows the mean of LAEPs across all listeners for each condition. A characteristic LAEP wave morphology was found for all masker types with a small positive deflection (P1), followed by a large negative deflection (N1) and a large positive deflection (P2). Fitted N1 and P2 amplitudes as a function of the target tone level are shown in Fig 8 and 9, respectively. Each panel shows the LAEPs of each masker type with both diotic (solid lines) and dichotic (dashed lines) target tones. As shown in Fig 10, both components showed an increase in amplitudes with increasing levels. Compared to N1, P2 amplitudes showed better goodness of fit for the power law function. In addition, N1 amplitudes showed more separation between diotic and dichotic conditions compared to P2 amplitudes.

**Fig 7:**
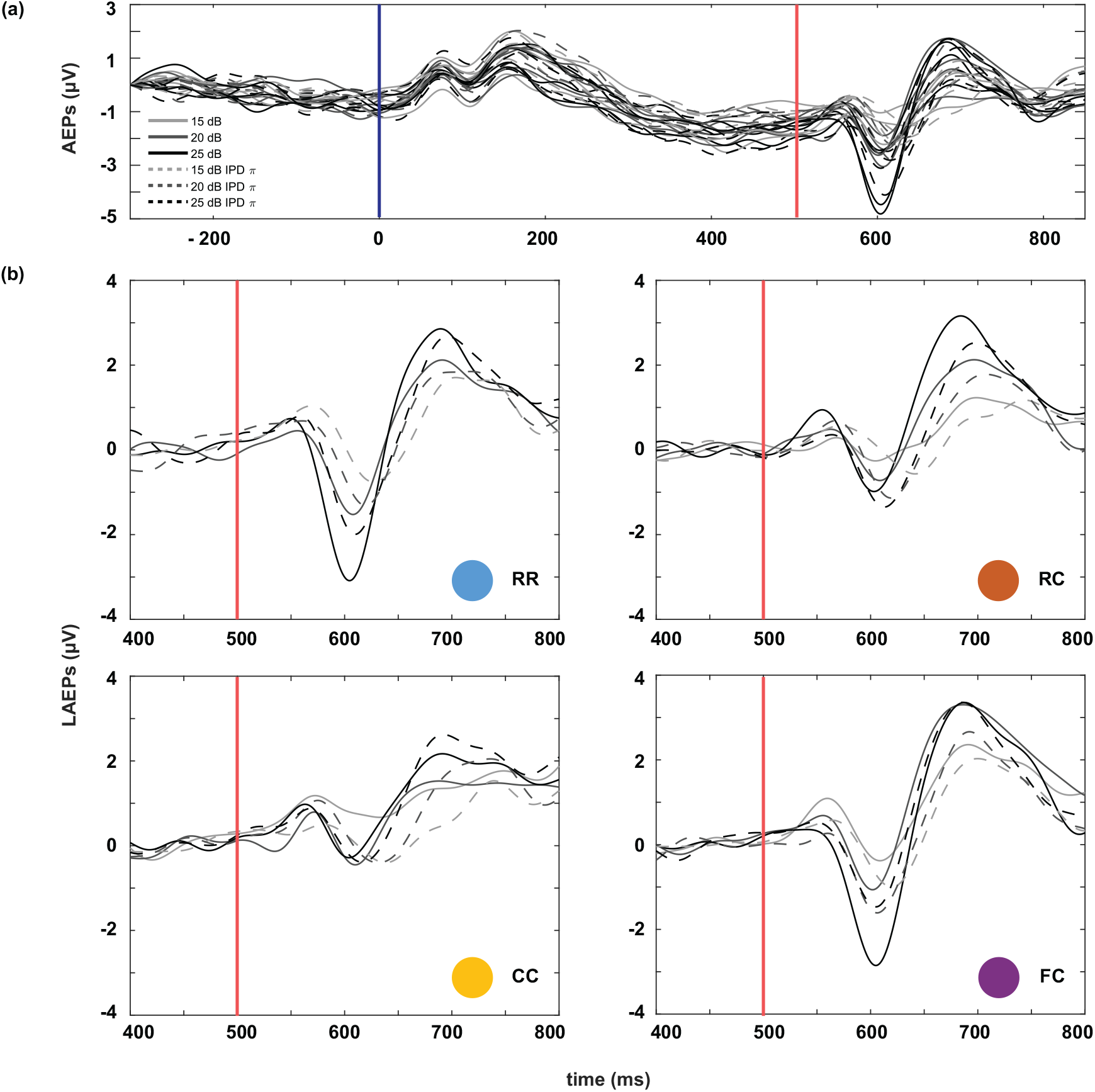
**Auditory evoked potentials (AEPs) averaged over all listeners. Masker onset is at t=0, target tone onset at t = 500 ms. Solid lines represent the four masker types with diotic target signals, and the dotted lines represent the four masker types with dichotic target signals. (b) Late auditory evoked potentials (LAEPs) to the target tone in the time interval ranging from 400 ms to 800 ms post masker onset**.

**Fig 8:**
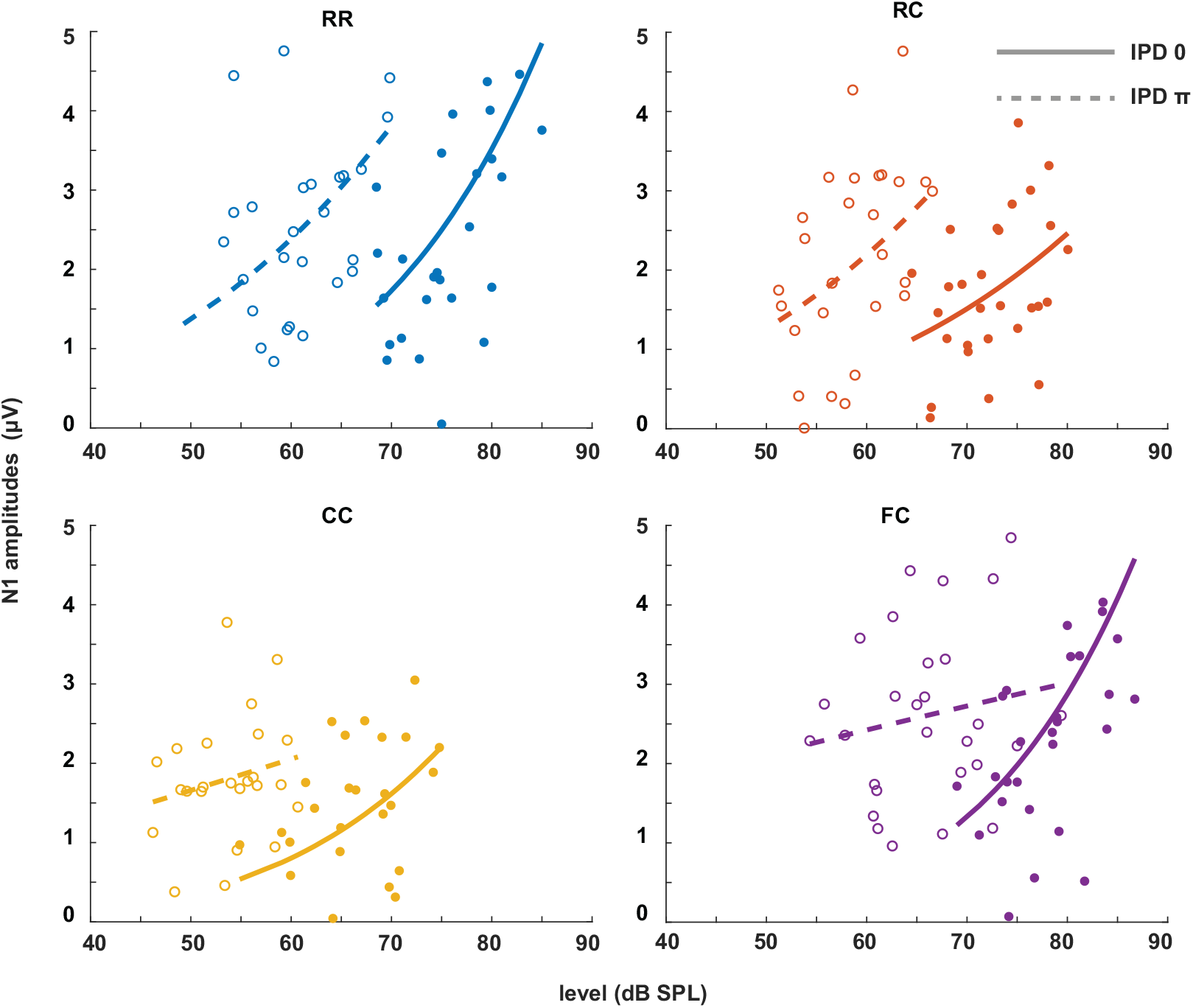
**N1 amplitudes as a function of target tone level at supra-threshold levels: + 15 dB, + 20 dB and + 25 dB. Individual data are plotted as single points. The data for each condition are fitted with a power function (line). Blue represents the RR condition, orange the RC condition, yellow the CC condition, and purple the FC condition. The solid lines represent the data of IPD 0 and the dotted line the data of IPD** *π*. **Foreach condition, the goodness of fit (***R*^2^**) with IPD of 0 was: RR(***R*^2^**=0.2776), RC(***R*^2^**=0.1657), CC(***R*^2^**=0.2179), FC(***R*^2^**=0.2893). The goodness of fit with IPD of** *π* **was: RR(***R*^2^**=0.2075), RC(***R*^2^**=0.1390), CC(***R*^2^**=0.0469), FC(***R*^2^**=0.0288)**.

**Fig 9:**
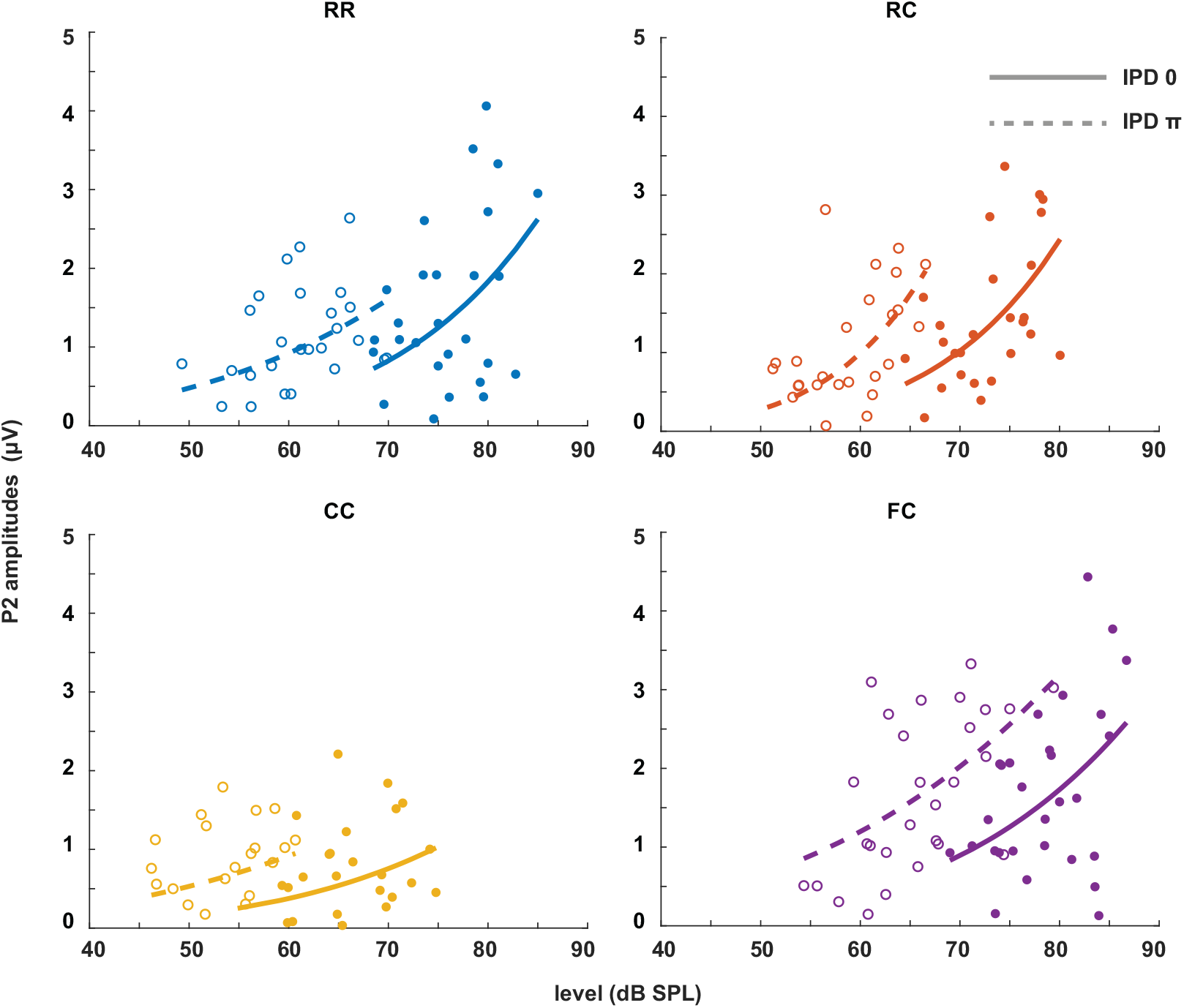
**P2 amplitudes as a function of target tone level at supra-threshold levels: + 15 dB, + 20 dB and + 25 dB. Individual data are plotted as single points. The data for each condition are fitted with a power function (line). Blue represents the RR condition, orange the RC condition, yellow the CC condition, and purple the FC condition. The solid lines represent the data of IPD 0 and the dotted line the data of IPD** *π*. **For each condition, the goodness of fit with IPD of 0 was: RR(***R*^2^**=0.1820), RC***R*^2^**=(0.3253), CC(***R*^2^**=0.0970), FC(***R*^2^**=0.1646). The goodness of fit with IPD of** *π* **was: RR(***R*^2^**=0.1847), RC(***R*^2^**=0.3224), CC(***R*^2^**=0.0601), FC(***R*^2^**=0.3161)**.

**Fig 10:**
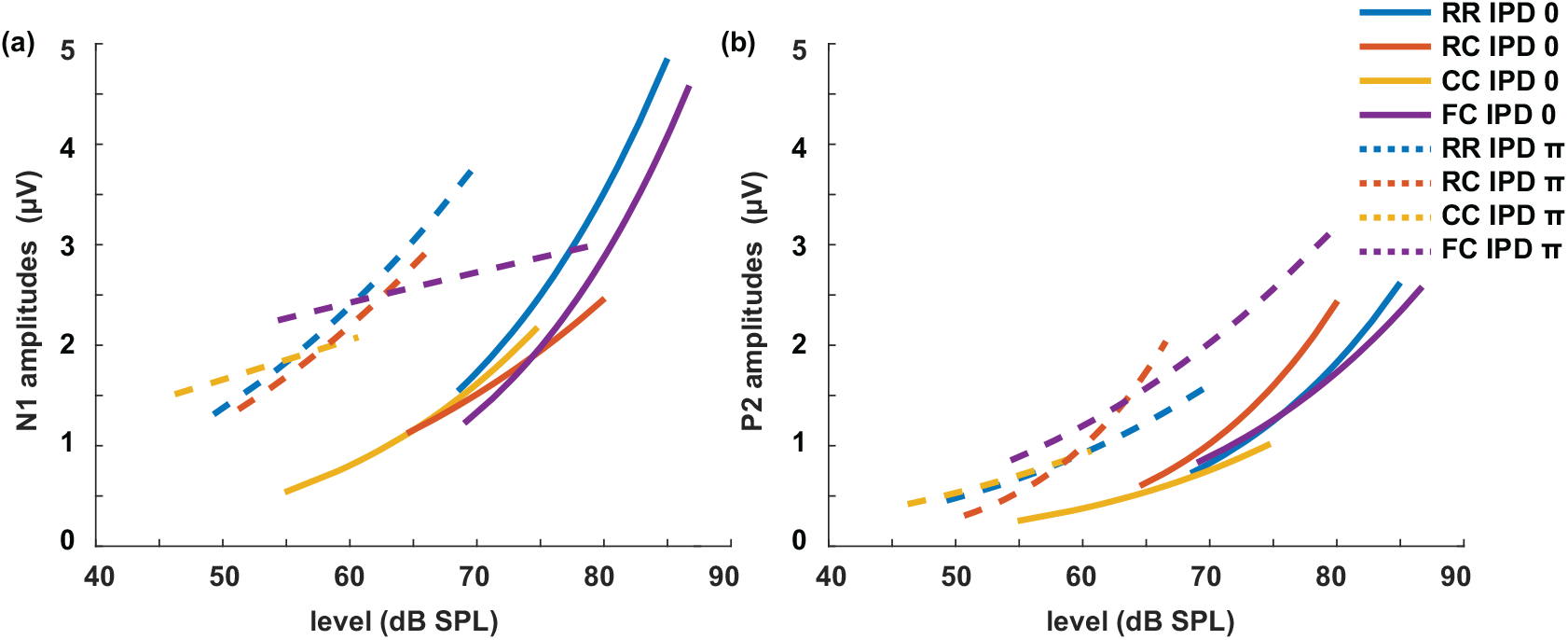
**The plots of the LAEPs with a function of target tone level. The data of N1 (left) and P2 (right) is fitted with the power function and plotted with the line. Blue corresponds to RR condition, orange to RC, yellow to CC, and purple to FC condition. Solid lines represent the data of IPD 0, and dotted lines represent the data of IPD** *π*.

## 4. Discussion

### 4.1. Effect of preceding maskers on CMR and BMLD

The results of the first experiment (Fig 2) showed that comodulation, IPD, and the preceding masker all influence the amount of masking release. In both diotic and dichotic conditions, the effect of preceding maskers on CMR was similar, as shown in Fig 2b. The amount of CMR (Fig 2b) was highest for the condition where the masker was comodulated for the whole duration of the interval (*CC*). CMR was reduced if the preceding masker had uncorrelated intensity fluctuations across frequency (*RC*). CMR was negative in the condition where comodulation of the preceding masker only spanned the FBs (*FC*). In a previous study by Grose et al. (2009), when the target tone was preceded and followed by maskers (temporal fringe), similar results were found. They interpreted the results in the light of the formation of a stream facilitated by the preceding masker. Even though the stimuli in the present study had no following masker after the offset of the target tone, the thresholds were in line with the results in (Grose et al., 2009) with both preceding and following maskers in a CMR paradigm. This suggests that the preceding masker plays a strong role in inducing auditory streams, which may impede the following stream formation by comodulation. This is also consistent with studies where the reduction of CMR by preceding or following stream formation was suggested as a high-level auditory processing (Dau et al., 2005, 2009). In addition,CMR was significantly reduced in dichotic conditions (e.g., *CC*_0_ vs. *CC*_*π*_). This is also in line with previous studies by (Schooneveldt and Moore, 1989; Cohen and Schubert, 1991; Ernst and Verhey, 2006; Epp and Verhey, 2009).

While the effect of preceding maskers on CMR was strong, its effect on BMLD was less pronounced. The amount of BMLD (Fig 2c) was similar across conditions and only showed a significant difference between the RR and the CC condition. The BMLD in the *CC* condition was lower compared to the *RR* condition. A potential reason for this reduced BMLD could be that the overall improvement of the target signal by comodulation and IPD reached a maximum. A similar phenomenon was observed in Epp and Verhey (2009) where listeners with a high BMLD showed slightly reduced CMR. Interestingly, the *FC* condition showed high individual variability in detection thresholds when the tone was presented with an IPD of *π*. From additional linear regression analysis (Fig 13), the *FC* condition in dichotic condition showed positive relation between CMR and BMLD. For listeners with low CMR and BMLD, the FBs and the CB might have been separated into different objects by comodulated FBs in the preceding masker. This may induce difficulties in separating the center masker from the tone due to its tone-like perceptual quality, especially when the target tone is presented at levels as low as 45 dB. For listeners with high CMR and BMLD, if they focused on the IPD cue, spatial information effectively separated the target tone from the noise and the noise components with no interaural disparity were grouped into one stream. However, this needs to be further investigated how the individual variability occurs.

We hypothesized that the preceding maskers would affect both CMR and BMLD if the effect of auditory object- or stream formation on masking release is due to higher-level auditory processing where prior knowledge affects sound perception. As previously mentioned, physiological evidence shows that neural correlates of comodulation processing can be found as early as the CN level (Pressnitzer et al., 2001; Neuert et al., 2004), while there is broad consensus that binaural information is processed at the level of the IC (e.g. Shackleton et al., 2003, 2005; Zohar et al., 2011). Under the assumption of bottom-up processing of the masked signal along the auditory pathway, beneficial auditory cues enhance the internal representation of the target signal at the brainstem level (CN, IC), inducing masking release. Hence, if the effect of the preceding maskers is the additional high-level auditory processing, on top of the brainstem level processing, this would affect the combined CMR and BMLD. In this study, the data suggest that BMLD is hardly affected by preceding maskers, while CMR varies strongly dependent on the type of preceding masker. This is not in agreement with the interpretation that the effect of preceding maskers is the result of high-level auditory processing (e.g., temporal integration).

**Table 1:** Linear regression summary for all conditions. *a* is CMR *b* is BMLD.

| | | $y = a * x + b$ | | |
| --- | --- | --- | --- | --- |
| | | $a$ | $b$ | $p$-values |
| diotic | RC | -0.04 | 14.08 | 0.886 |
|  | CC | -0.56 | 18.05 | 0.044* |
|  | FC | -0.56 | 11.18 | 0.338 |
| dichotic | RC | 0.01 | 13.95 | 0.968 |
|  | CC | 0.13 | 11.75 | 0.610 |
|  | FC | 0.83 | 18.13 | 0.006* |

A possible explanation is the top-down processing where prior knowledge about the sound is used to influence the processing of sensory information at the low-level (Asilador and Llano, 2021). In this scenario, the auditory system uses accumulated information of incoming sound, which can be understood as adaptation at a “system-level”. This adaptation at the cortical level could affect auditory processing at the brainstem. Such an auditory efferent system from the auditory cortex to the CN could explain the effect of preceding maskers on CMR but not BMLD (Terreros and Delano, 2015). However, the neural correlates for the top-down modulation arising from the preceding maskers are unknown.

Lastly, one might also speculate about the role of adaptation processes at the peripheral level in the effect of preceding maskers. Similar to the paradigm used in this study, various psychophysical and neural phenomena have shown the influence of preceding signals on the following target tone perception, termed as “auditory enhancement” (e.g. Nelson and Young, 2010; Kreft et al., 2018). In these studies, the preceding maskers were broadband noise with a spectral notch around the target signal. The presence of a spectral gap around the signal frequency in the preceding masker enhanced the target detection. The underlying mechanism of “auditory enhancement” has been attributed to the adaptation at both low- and high-level auditory processing. For supporting the adaptation at low-level auditory processing, Kreft et al. (2018) suggested that olivocochlear efferents may induce the adaptation effect in a longer time scale than the auditory nerve fibers (Guinan Jr, 2006). If this is the case, how modulation patterns (e.g., *RR, RC, CC*, and *FC*) result in different degrees of CMR reduction is in question. Based on a physiological study where modulation pattern encoding was found at the CN level, the connectivity between the CN and the medial olivocochlear (MOC) efferents may play a role (Pressnitzer et al., 2001; Oertel et al., 2011). However, further psychoacoustic and physiological studies are needed to develop current ideas.

### 4.2. Benefit of CMR and BMLD at supra-threshold levels

The results of the second experiment (Fig 3) showed that the intensity JND was inversely proportional to the physical sound level. This is consistent with data from the literature for pure tones in quiet (e.g. Ozimek and Zwislocki, 1996) where the intensity JND decreased according to the power function of sensation level. Expression of the JND on a relative scale to the reference level (10*log*(Δ*L/L*)) showed independence of the JND on the masker type (*RR, RC, CC, FC*) and the IPD (0, *π*). This means that, for a given target tone level, regardless of the difference in masked thresholds, the intensity JND on a relative scale was the same. Such level dependency of JND is interesting in terms of the level above the masked threshold or the supra-threshold level. Between two conditions, the level above masked threshold can differ by up to 25 dB at a given target tone level (*FC*_0_ vs. *CC*_*π*_). While the target tone level of 70 dB SPL is just above the threshold for the FC masker and well above the threshold for the CC masker. Still, for both cases, the same relative amount of intensity increment was required for the discrimination.

It is often assumed that the neural encoding of sound intensity is implemented by spike rate (Cai et al., 2009; Micheyl et al., 2013b). However, auditory nerve fibers (ANFs) usually saturate above certain sound levels (Bruce et al., 2018). Therefore, if the intensity JND measures are the result of rate-based encoding, an additional mechanism must exist to combine information across ANFs (Viemeister, 1988). Several studies have suggested that the auditory cortex plays such a role in intensity discrimination (Dykstra et al., 2012; Micheyl et al., 2013a). We propose that such a mechanism could also be located at the level of the CN. Physiological studies found neural correlates of CMR where neural activity was affected by comodulation (e.g. Nelken et al., 1999; Pressnitzer et al., 2001; Neuert et al., 2004). One might think that at a given stimulus level, the neural activity is higher in conditions with a masking release compared to a condition without a masking release. In this case, it seems plausible that the internal representation of the tone rather than the physical target tone level is relevant for sound perception. However, as the intensity JND is the same for the same target tone level regardless of the amount of masking release, our results indicate that the physical target tone level is encoded and preserved, in addition to the enhanced neural representation at thresholds as an internal signal-to-noise ratio (iSNR). For the intensity encoding at the level of CN, small cells showed preserved intensity encoding of the target tone in the presence of the noise (Hockley et al., 2022). These cells displayed a unique rate-level function where the spike rate increases without saturation with increasing levels up to 90 dB SPL (Hockley et al., 2022). This could be a possible mechanism of the intensity coding in masking release conditions.

### 4.3. Estimation of salience with the intensity JND

We estimated salience rating based on the intensity JND measurements. The salience rating at the threshold was set to on arbitrarily. This was based on the idea that the detection of a signal by the auditory system is possible once the internal representation of that signal exceeds a critical iSNR. Based on linear signal theory approach, any addition of signal energy above the detection threshold should increase the iSNR proportionally with the increase of the signal intensity, resulting in enhanced salience (e.g., Epp and Verhey, 2009; Egger et al., 2019). The data from the present study, however, is not in line with this hypothesis. As shown in Fig 6, the salience increases as a function of the target tone level, but each condition shows different slopes rather than constant slopes. This means that the change in salience is dependent on the physical target tone level rather than the iSNR. At higher intensities, the estimated salience measures converge, indicating the vanishing effect of the beneficial cues leading to CMR and BMLD. This suggests that beneficial cues for target detection might not be used by the auditory system at natural conversational levels.

From a physiological point of view, this interpretation would imply that the physical target tone level needs to be encoded, affecting the neural representation of the signal enhanced by the processing of comodulation and IPD. This also clearly outlines a shortcoming of the simplified model by Epp and Verhey (2009), which does not reflect any nonlinearity that would explain this behavioral outcome. Thus, further studies are needed to enable us to extend this argument towards more complex signals like speech. Furthermore, it should be highlighted that the estimated salience in the present study and in the study by Egger et al. (2019) likely reflect different aspects of perception. Egger et al. (2019) suggested that some listeners might have used a partial loudness cue to assess the salience of the presented target tone. This is consistent with the present study in terms of the dependence on the physical target tone level rather than the level above the masked threshold. However, with existing loudness model, the relation between the salience and loudness growth as a function of the target tone level in masking release conditions is unclear.

### 4.4. LAEPs and intensity JNDs

In previous studies, the P2 component of the LAEP was suggested to be correlated with the supra-threshold levels of the masked tone (Epp et al., 2013). They showed that P2 amplitudes were proportional to the amount of masking release, CMR, and BMLD. Their results were based on measurement of the LAEP at fixed target tone levels in the absence and presence of comodulation and IPD. In a follow-up study by Egger et al. (2019), they measured P2 amplitudes at the same supra-threshold levels that were adjusted individually for all listeners. They found that the P2 amplitudes were similar at the same level above threshold across conditions. They noted that N1 amplitudes were correlated with the amount of BMLD. They also measured the salience of the target tone with the continuous scaling method. However, they could not find the correlation between the P2 amplitudes and the salience ratings. In the present study, we estimated how P2 amplitudes grow as a function of the target tone level. As shown in Fig 10, N1 amplitudes showed more prominent difference between diotic and dichotic conditions than P2 amplitudes. However, the *FC*_*π*_ showed a little correlation with target tone levels than other conditions. If N1 amplitudes reflect BMLD processing at the IC level, this might suggest an additional higher-level BMLD processing. On the contrary, P2 amplitudes were proportional to the target tone level in all conditions, and showed higher goodness of fit than N1 amplitudes. P2 amplitudes were larger in dichotic conditions than diotic conditions, reflecting enhanced salience. Between conditions with the same IPD, difference was marginal compared to estimated salience behaviorally. In addition, with a hypothesis that the intensity JND is encoded with spike rate, we estimated LAEPs and the intensity JND measures (Fig 11). We estimated both LAEPs and the intensity JND from the fitted function of LAEPs and intensity JNDs. We used individual supra-threshold levels of all conditions from fifteen listeners as an input to the fitted functions. The amplitudes of LAEPs were inversely correlated with the intensity JND measures and re-scaled JND. P2 amplitudes showed a steeper increase as the intensity JND decreases. With re-scaled JND, which showed better correlation with the target the level (Fig 5), P2 amplitudes across conditions with same IPD were less diverted from each other compared to the intensity JND measures. P2 amplitudes had a linear relationship to re-scaled JND measures.

**Fig 11:**
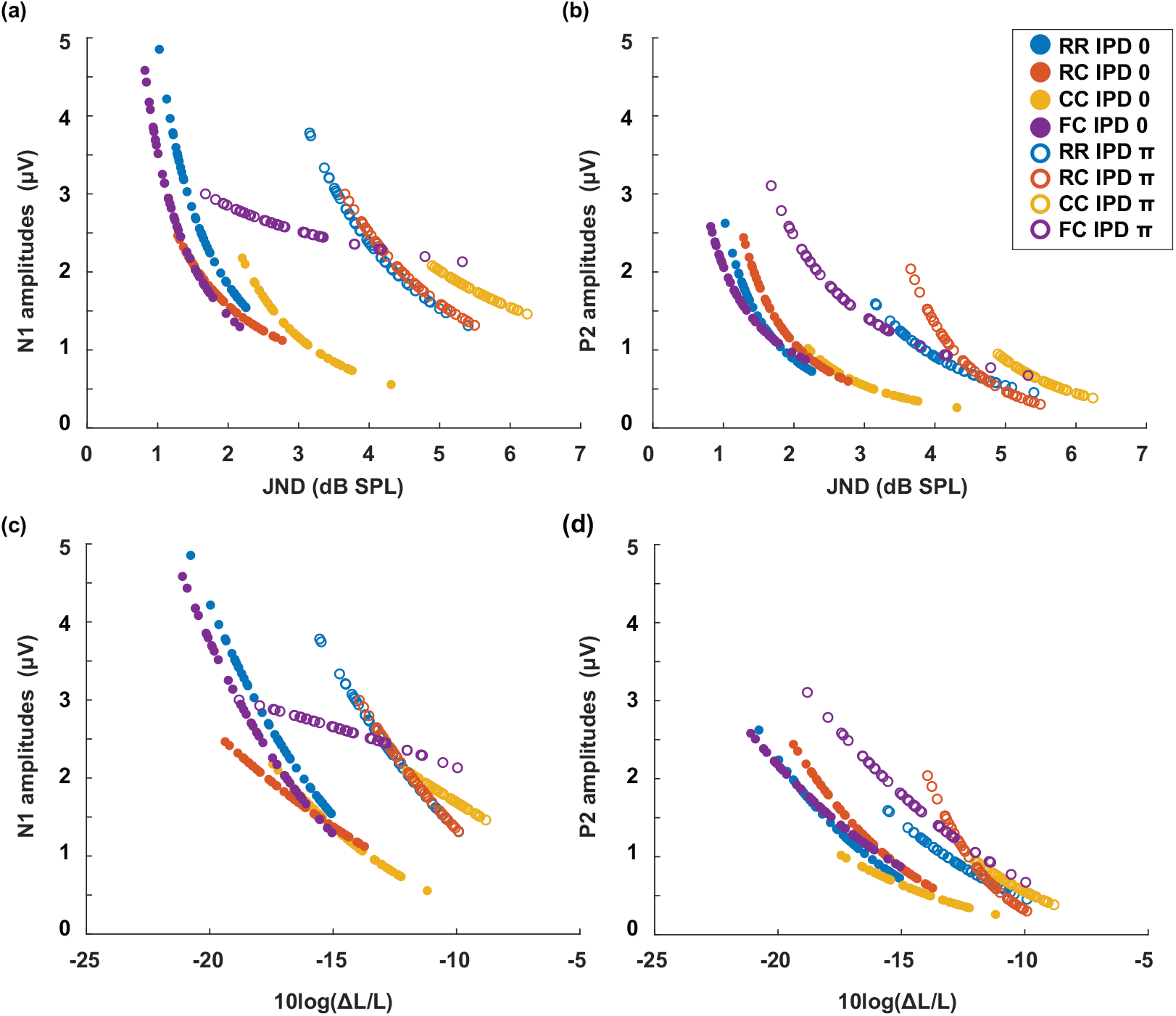
**LAEPs as a function of re-scaled intensity JNDs (**Δ*L/L***). The data of N1 (left) and P2 (right) are fitted with a power function. The blue line represents the** *RR* **condition, the orange line the** *RC* **condition, the yellow line the** *CC* **condition, and the purple line the** *FC* **condition. The solid line represents the data for IPD of 0 and dotted lines the data for IPD of** *π*.

### 4.5. LAEPs and salience

To investigate if P2 amplitudes could be a neural measure for salience, we estimated the salience at the supra-threshold levels individually (+15 dB, + 20 dB, + 25 dB). Fig 12 shows the relation between the estimated salience and the amplitudes of N1 and P2. Although P2 amplitudes were more correlated with estimated salience than N1 amplitudes, two conditions showed deviating patterns in dichotic conditions (e.g., *RC*_*π*_, *FC*_*π*_). Here, we assumed that the internal representation of the target tone in noise arises from serial auditory processing, and the resulting iSNR is reflected in P2 amplitudes. In the first experiment, CMR and BMLD showed non-linearity, such as reduced CMR in dichotic conditions and a strong correlation between CMR and BMLD in the FC dichotic conditions. In the second experiment, the intensity JND showed high variance in low target level compared to the re-scaled JND measures. As estimated salience was based on the intensity JND measures, this might have affected the accuracy of the salience rating. Therefore, salience estimation based on re-scaled JND measures may produce a better prediction. However, the method for translating re-scaled JND measures to salience measures needs to be further investigated. In the third experiment, N1 amplitudes were not correlated with the audibility, or BMLD processing, in the *FC*_*π*_ condition. This also suggests a possible higher-order auditory processing that may play a role in shaping neural responses. As the neural mechanisms underlying such non-linearity is unclear, further physiological evidence is needed to make a clear conclusion on how much extent P2 amplitudes can reflect the auditory processing stages and predict the salience. If additional high-level auditory processing is involved in combining CMR and BMLD together with temporal integration, AEPs that elicited later than P2 (e.g., P300) might provide more insights on the feasibility of electrophysiological measures for the salience.

**Fig 12:**
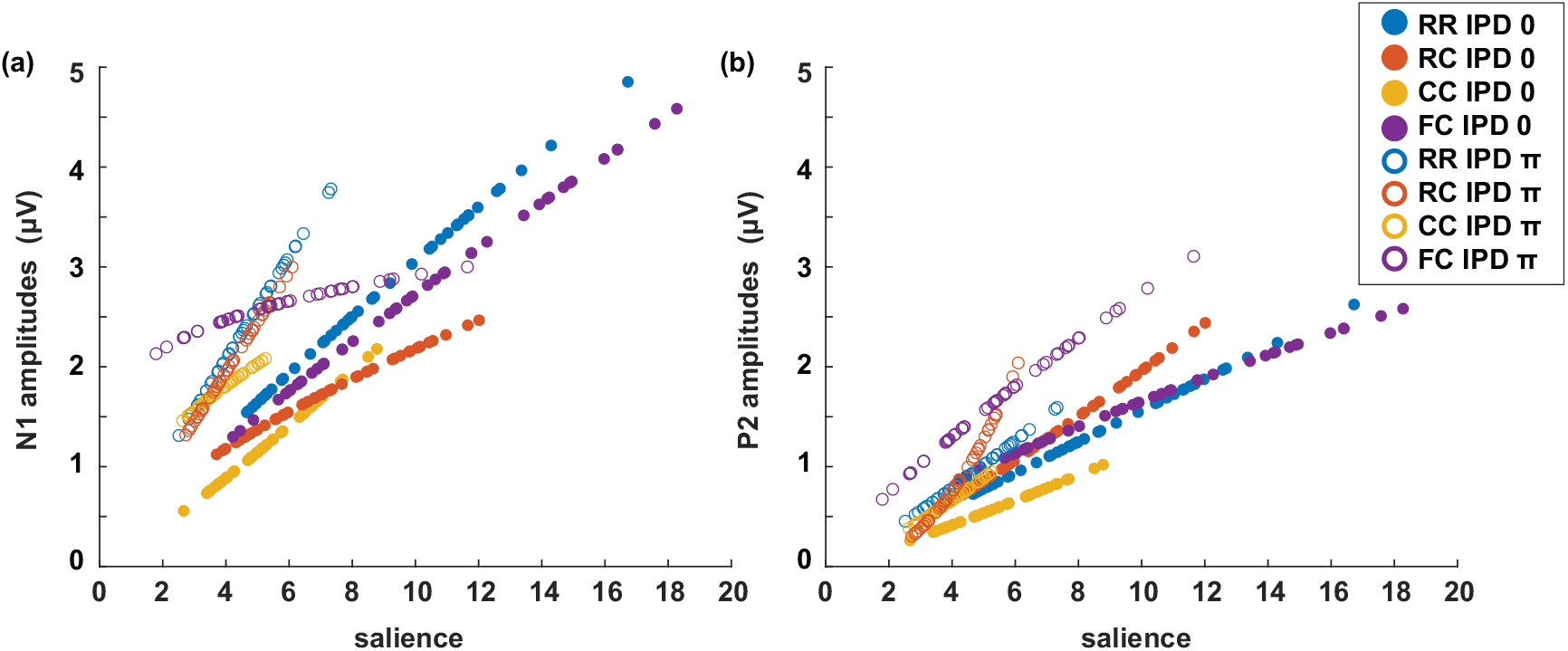
Estimated salience correlated with a) the N1 amplitude of the LAEP, and b) the P2 amplitude of the LAEP.

**Fig 13:**
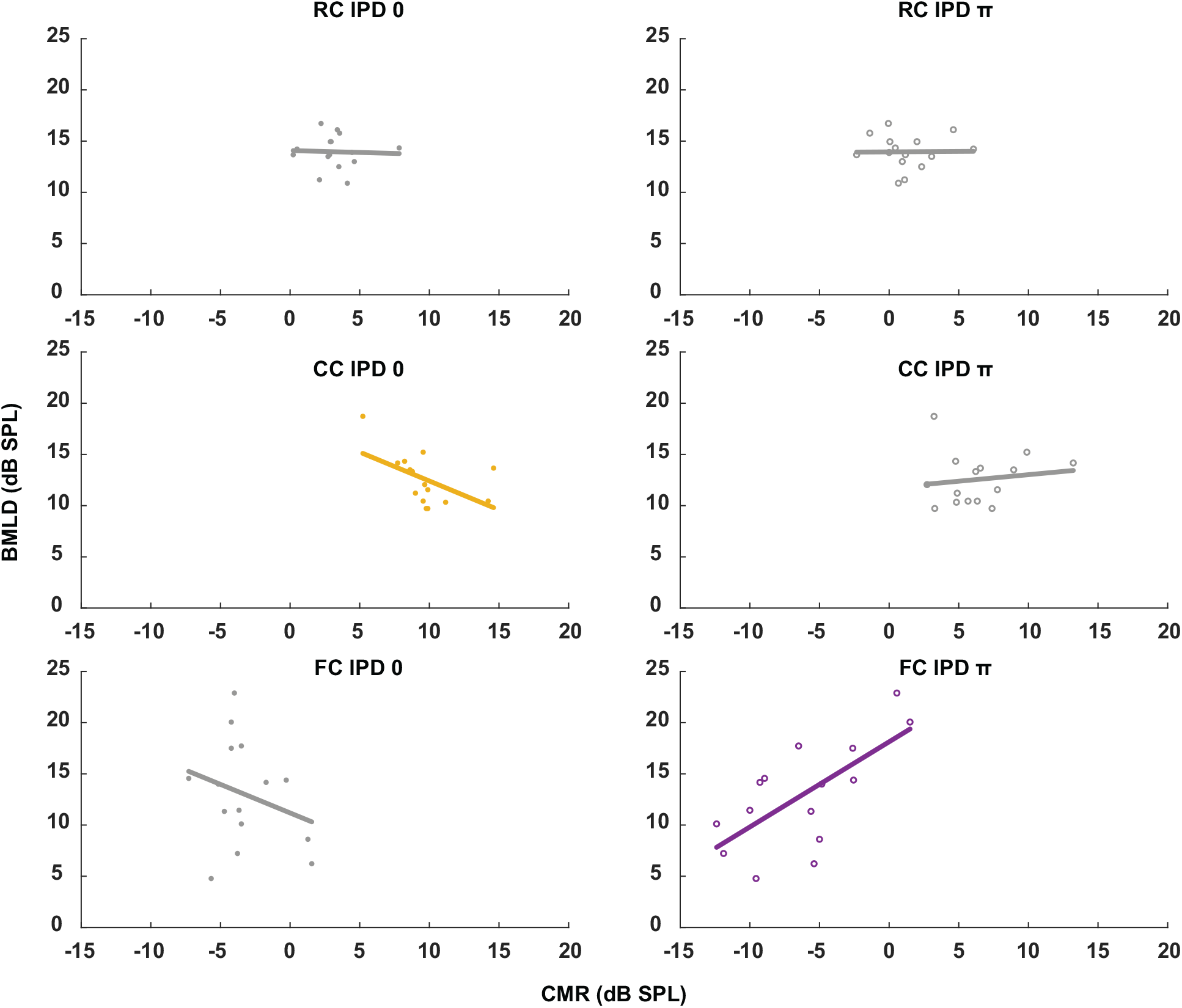
**Linear regression analysis between CMR and BMLD for all stimulus conditions. Only the conditions with significant p-values were plotted with color (see Table 1)**.

## 5. Conclusion

In this study, we investigated the detection and discrimination of masked tones in masking release conditions. Auditory cues such as comodulation and IPD, and the preceding masker, could enhance the detection performance. On the other hand, at supra-threshold levels, the discrimination performance was highly dependent on the physical target tone level. Regardless of the masking release conditions, the intensity JND measures were correlated with the target tone level. Furthermore, the estimated salience was higher in conditions with lower detection thresholds. At the high level, however, estimated salience converged to the same value across conditions. Lastly, the P2 amplitudes were more correlated with the behavioral measure of the salience than the L1 amplitudes.

## Acknowledgments

We would like to thank Viktorija Ratkute for support with the data collection.

